# Correcting Modification-Mediated Errors in Nanopore Sequencing by Nucleotide Demodification and in silico Correction

**DOI:** 10.1101/2022.05.20.492776

**Authors:** Chien-Shun Chiou, Bo-Han Chen, You-Wun Wang, Nang-Ting Kuo, Chih-Hsiang Chang, Yao-Ting Huang

## Abstract

The accuracy of Oxford Nanopore Technology (ONT) sequencing has significantly improved thanks to new flowcells, sequencing kits, and basecalling algorithms. However, novel modifications untrained in the basecalling models can seriously reduce the quality. This paper reports a set of ONT-sequenced genomes with unexpected low quality (∼Q30) due to extensive new modifications. Demodification by whole-genome amplification (WGA) significantly improved the quality of all genomes (∼Q50-60) while losing the epigenome. We developed a computational method, Modpolish, for correcting modification-mediated errors without WGA. Modpolish produced high-quality genomes and uncovered the underlying modification motifs without loss of epigenome. Our results suggested that novel modifications are prone to ONT errors, which are correctable by WGA or Modpolish without additional short-read sequencing.

## Background

The Oxford Nanopore Technology (ONT) is a popular long-read sequencing platform that enables real-time sequencing for point-of-care medical applications, such as the diagnosis of infectious and newborn diseases within hospitals [1, 2]. Despite its great potential and popularity, the accuracy of ONT was inferior to those of other platforms (e.g., Illumina and PacBio HiFi). Recently, the quality of ONT sequencing has significantly improved thanks to new flowcells (e.g., R10.4), sequencing kits (e.g., Kit 14), and basecalling algorithms (e.g., Bonito). For example, by using the R10.4 flowcells, near-perfect microbial genomes from isolates or metagenomes can be reconstructed by ONT-only sequencing without short-read polishing [3].

However, because the throughput of R10.4 is much lower due to slower sequencing speed, most projects rely on the R9.4 flowcells for higher yield. Although the upcoming sequencing kit will further enhance the accuracy (e.g., Kit 14), postassembly genome polishing is still compulsory for removing ONT systematic errors regardless of the flowcell or kit versions. Systematic errors are recurrent basecalling errors at the same locus, which are not correctable by the consensus of read pileups (e.g., Racon) [4]. Homopolymer errors (i.e., indels) were the primary source of ONT systematic errors. Thanks to several machine-learning algorithms, these errors have been significantly reduced by read-based (e.g., Medaka) or reference-based (e.g., Homopolish) polishing methods [5]. These algorithmic advances have produced high-quality ONT genomes sufficient for downstream analysis (e.g., >Q50) [3, 6].

Unfortunately, the ONT signals are ultra-sensitive to various modifications (e.g., 5mC, 6mA). More than 17 and 160 modification types have been found in DNA and RNA, respectively, and the number is still growing (e.g., DNA adducts, N4-acetyldeoxycytosine) [7, 8]. These modifications disturb the electrical current and result in unfixable systematic errors [9]. Note that these modification-mediated errors cannot be eliminated by new flowcells and sequencing kits (e.g., R10.4 and Kit 14) which aim to reduce homopolymer errors. Furthermore, existing basecalling and polishing algorithms (e.g., Guppy and Medaka) were trained for capturing only a few modifications (e.g., 5mC, 5hmc, 6mA). Consequently, the quality of ONT sequencing is unreliable when novel modifications extensively edit the genome.

This paper presents a set of unexpected low-quality genomes due to extensive novel modifications. We show that the removal of modifications by whole-genome amplification (WGA) significantly improves the quality of all genomes. A novel computational method is developed for correcting these modification errors without WGA.

## Results

### Unusual low-quality of ONT genomes due to extensive modifications

We sequenced 12 microbial strains of *Listeria monocytogenes* using Illumina and ONT (∼200-990Mbp) (Figure 1(a), Supplementary Tables S1 and S2). The ONT reads were assembled into genomes with sequencing errors further polished by the-state-of-the-art tools (Supplementary Table S3, see Methods). The Illumina and ONT reads were hybrid assembled for evaluation purposes (Supplementary Table S4). When compared with the Illumina/ONT hybrid assemblies (Figure 1(b)), seven ONT-only genomes exhibited high quality (HQ) ranging from Q47 to Q60 (e.g., R19-2905 and R20-0088). However, five isolates (R20-0026, R20-0030, R20-0127, R20-0148, and R20-0150) showed unexpectedly low quality (LQ) varying from Q27 to Q34. The accuracy of these five LQ genomes remained unimproved after replicated ONT sequencing (data not shown). Further investigation of the five LQ genomes revealed excessive amounts of mismatch errors (1,228-5,780) compared with the seven HQ ones (3-36 mismatches) (Figure 1(c)). Homopolymer errors (i.e., indels) were not the source of inferior quality (7-306, Supplementary Table S5).

**Figure 1.**
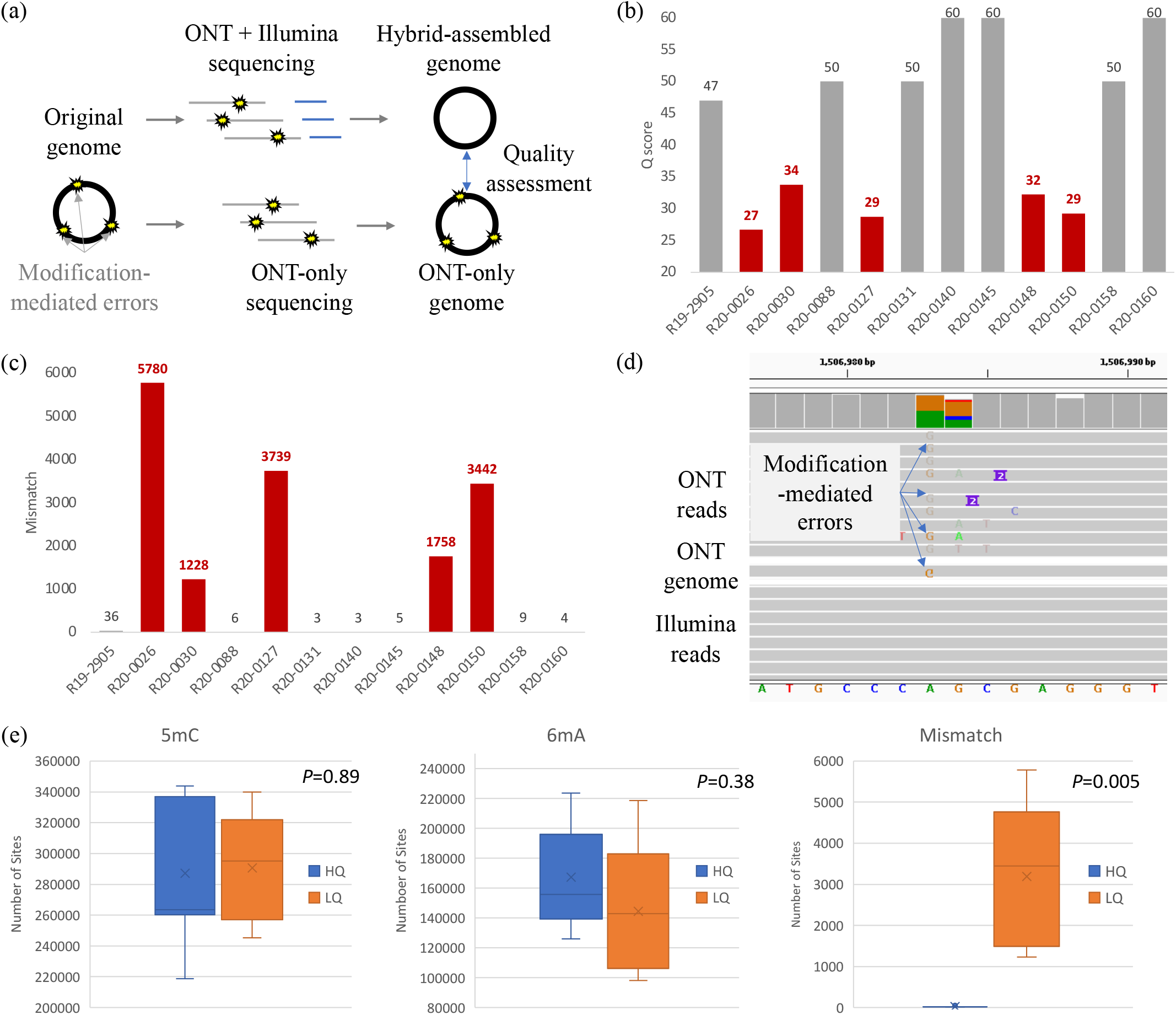
Quality comparison of 12 microbial strains using ONT-only and ONT/Illumina hybrid sequencing. (a) Workflow of ONT-only and ONT/Illumina hybrid assembly; (b) Q scores; (c) number of mismatches; (d) comparison of ONT and Illumina reads by IGV; (e) numbers of 5mC, 6mA, and mismatches between HQ/LQ strains.

Manual inspection revealed that these mismatches were ONT basecalling errors uncorrected after genome polishing (Figure 1(d) and Supplementary Figure S1). As mismatch errors in ONT are mainly due to epigenetic modifications, we computed the frequency of well-known methylation in these isolates (see Method and Supplementary Table S6). In terms of 5-methylcytosine (5mC), the numbers of modified loci in the five LQ genomes (∼240-340k) were not significantly higher than those in the HQ ones (210-345k, *P*=0.89, Figure 1(e)). Similarly, the numbers of N^6^-methyladenine (6mA) modifications also showed no significant difference between the LQ and HQ groups (98-218k v.s. 126-223k, *P*=0.34, Figure 1(f)). Because the numbers of mismatch errors in LQ genomes are significantly higher than those of HQ ones (P=0.005, Figure 1(g)), we suspected ONT basecalling algorithms failed to distinguish the novel modifications in the LQ isolates.

### High-quality ONT genomes by WGA demodification

We removed the modifications in all microbial samples by WGA (Figure 2(a)), which randomly amplifies the genome fragments without retaining any epigenetic modification (see Methods). The WGA-demodified samples were sequenced by ONT, assembled into chromosomes, and compared with the Illumina/ONT hybrid genomes (Figure 2(a), Supplementary Tables S7 andS8). The five LQ genomes after WGA exhibited significantly higher quality than those without demodifications (e.g., Q27 to Q53 in R20-0026) (Figure 2(b), Supplementary Table S9). In particular, the amounts of mismatch errors significantly reduced after demodification (e.g., 5,780 to 16 in R20-0026) (Figure 2(c)). Consequently, the unexpected low quality of ONT was due to excessive novel modifications untrained in their basecalling model. The demodification by WGA can produce high-quality ONT genomes without the need for Illumina short reads.

**Figure 2.**
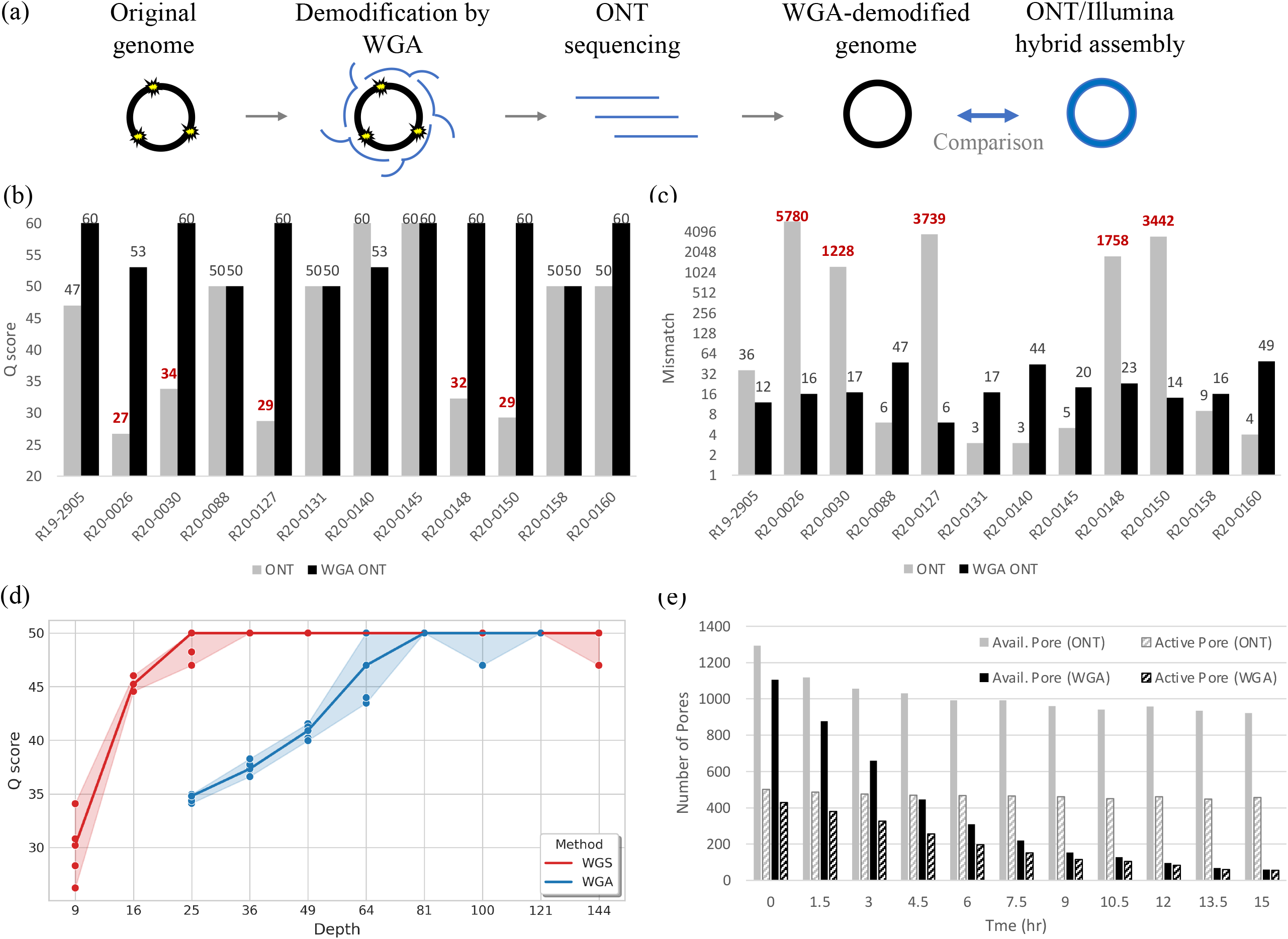
Quality improvement of ONT by WGA demodification. (a) Worflow of WGA-demodified ONT; (b) Q scores of the WGA-demodified and ONT-only genomes; (c) numbers of mismatches of the WGA-demodified and ONT-only genomes; (d) WGA genome quality with respect to sequencing depth; (e) numbers of active/available pores during WGA-demodified and ordinary ONT sequencing.

However, while WGA successfully erased these modifications, the sequencing cost increased by two factors. First, WGA required a higher sequencing depth (∼100x) for assembling a complete genome when compared with ordinary ONT sequencing (∼30x) (Figure 2(d) and Supplementary Figures S2-3). It was due to the uneven amplification of WGA, which led to non-uniform sequencing depth and a fragmented assembly at moderate coverage. Second, the WGA-demodified samples may reduce the ONT yields. We observed the numbers of available/active pores could sometimes decrease quickly (e.g., less than 100 pores after 12h) (Figure 2(e)), which was possibly owing to the hyperbranched structure unresolved after WGA. Consequently, the sequencing cost of WGA-demodified samples using ONT is much higher than ordinary sequencing.

### *in silico* correction of modification-mediated errors by Modpolish

We developed a novel computational method (called Modpolish) for correcting these modification-mediated errors without WGA and prior knowledge of the modifications. Modpolish identifies and corrects the modification-mediated errors by investigating basecalling quality, basecalling consistency, and evolutionary conservation (Figure 3(a), see Method). Briefly, because the ONT signals are disturbed by modifications, the basecalling quality is usually low, and the basecalled nucleotides are often inconsistent at the modified loci. In conjunction with the conservation degree measured by closely-related genomes, only the modified loci with ultra-high conservation will be corrected by Modpolish, avoiding false corrections of strain variations.

**Figure 3.**
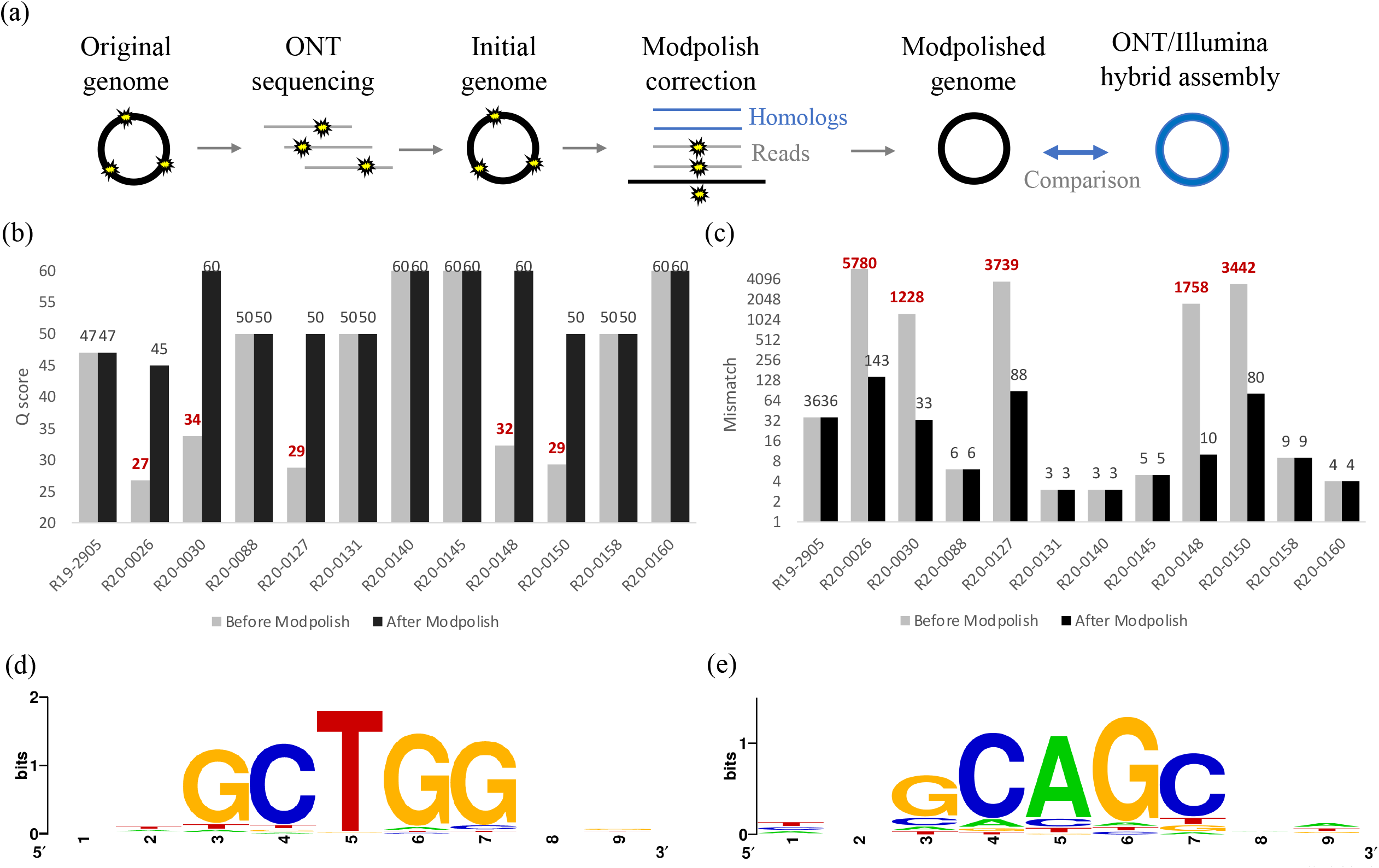
Correction of modification-mediated errors by Modpolish. (a) Workflow of Modpolish; (b) Q scores before and after Modpolish; (c) numbers of mismatchs before and after Modpolish; (d) the sequence motif of modification on ST1081; (e) the sequence motif of modifications on ST87.

We assessed the accuracy of Modpolish by comparing the quality of the ONT-only genomes (polished by Medaka/Homopolish) with those further polished by Modpolish. The results indicated that Modpolish significantly improved the genome quality of all LQ genomes (Figure 3(b), Supplementary Table S10). For instance, the quality of R20-0030 improved from Q34 to Q60, and the number of mismatches decreased from 1,228 to 33 (Figure (3(c)). We observed that the number of mismatches in R20-0026 reduced dramatically (i.e., from 5,780 to 143). However, the quality improvement (i.e., from Q27 to Q45) was slightly inferior to the others due to the 143 uncorrected mismatches. Note that no false corrections were made on the seven HQ genomes, implying the correction specificity of Modpolish is high.

The multilocus sequencing typing (MLST) indicated that R20-0026 belonged to the sequence type ST1081 and the remaining four LQ strains (i.e., R20-0030, R20-0127, R20-0148, R20-150) were ST87. Hence, we investigated whether an identical modification system extensively edited the genomes of these two lineages. Sequence analysis of the modified loci revealed that the modifications of ST1081 were on the GCTGG motif (Figure 3(d)). On the other hand, the modification sites of all ST87 strains centered on the GCAGC motif (Figure 3(e)). Therefore, two modification systems seem specific to each of the two lineages. In addition, while both motifs are not palindromic, their reverse complements (i.e., CCAGC, GCTGC) were also hotspots of modifications (Supplementary Figure S4). Because the mismatches frequently appeared on both strands at the same loci (Supplementary Figure S5), the unknown modification may symmetrically edit both strands. Although their underlying mechanisms remained unclear, the two systems extensively modified the genomes at specific motifs with high conservation within each lineage, leading to excessive amounts of basecalling yet correctable errors.

### Comparison of phylogeny reliability under extensive modifications

Because sequencing errors alter the genetic distances between strains, we assessed the reliability of phylogeny using ONT with or without modification-error removal. We reconstructed the core-genome MLST (cgMLST) phylogeny of the five LQ strains sequenced and assembled by four methods: ONT-only sequencing, WGA-demodified ONT, ONT with Modpolish, and hybrid ONT/Illumina sequencing (Figure 4(a)). The WGA-demodified genomes perfectly clustered with the ONT/Illumina hybrid for each strain in both clades (ST87 and ST1081). The ONT genomes corrected by Modpolish clustered with the hybrid and WGA-demodified genomes in both clades. But the genetic distance slightly deviated from them, especially in the ST1081 clade. The ONT-only genomes were phylogenetic distant from the others due to excessive amounts of modification-mediated errors.

**Figure 4.**
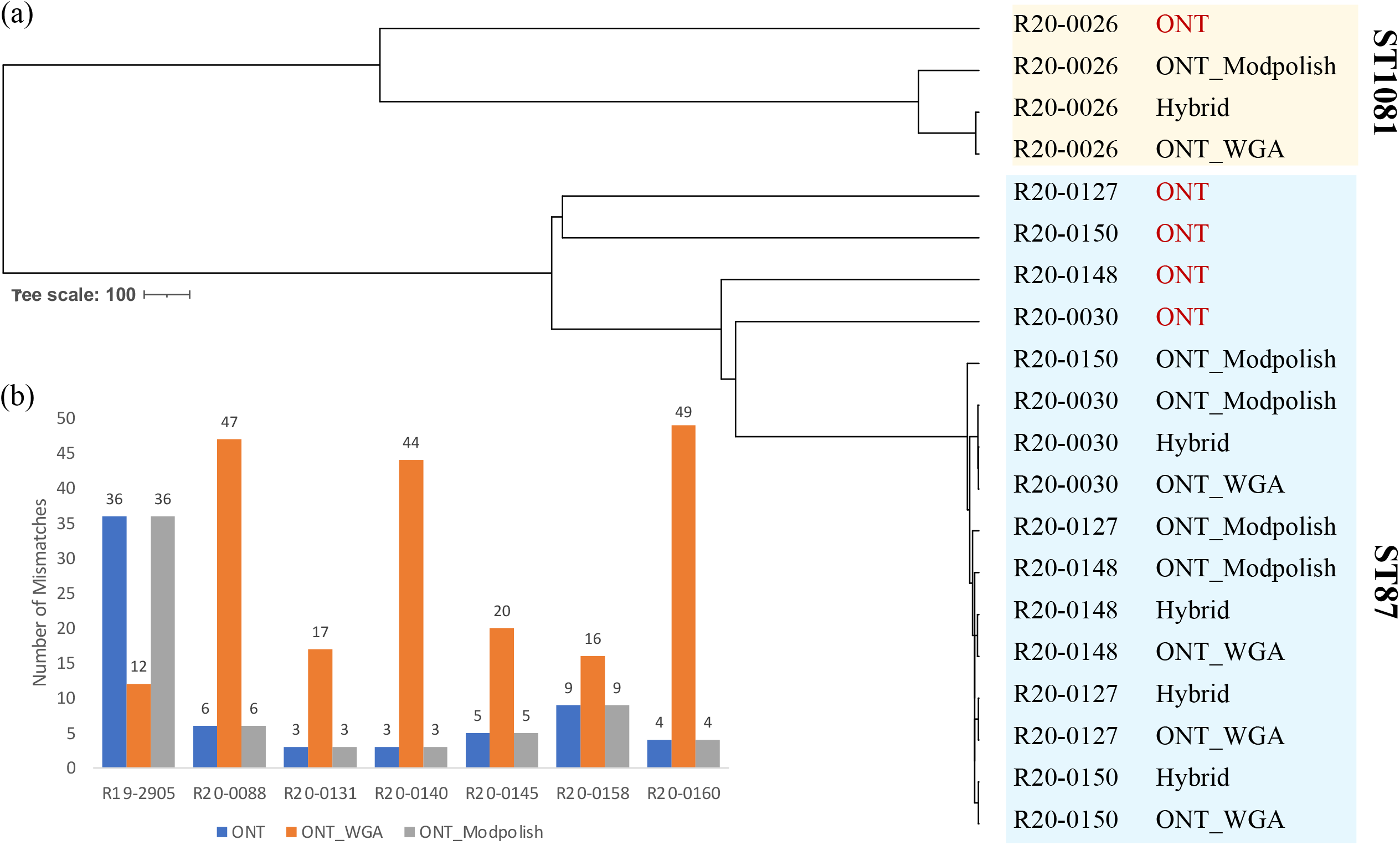
Comparison of phylogeny reliability of four methods. (a) The cgMLST phylogeny of the five LQ strains sequenced and assembled by four methods: ONT-only sequencing (ONT), WGA-demodified ONT (ONT_WGA), ONT with Modpolish (ONT_Modpolish), and hybrid ONT/Illumina sequencing (Hybrid_WGS); (b) the cgMLST distances of ONT, ONT_WGA, and ONT_Modpolish to the Hybrid_WGS assembled genomes.

When comparing each method in the seven HQ isolates, ONT with WGA was slightly worse than the original ONT and Modpolish in six strains (e.g., 47 v.s. 6 mismatches in R20-0088) (Figure 4(b)), except for the R19-2905 isolate (i.e., 12 v.s. 36 mismatches). These mismatches slightly increased the genetic distance to the others (Supplementary Figure S6). Nevertheless, phylogenetic analysis indicated that the genomes of all methods were perfectly clustered for each HQ strain (Supplementary Figure S7), implying the number of mismatches is less than that of strain variations. Consequently, all methods can produce reliable phylogeny when free of novel modifications. But when new modifications extensively edit the genome, only ONT with WGA or Modpolish can provide sufficient typing accuracy without additional Illumina sequencing.

## Discussion

This paper presented a set of unexpected low-quality ONT genomes due to extensive modifications untrained in the basecalling models. Demodification by WGA successfully improved the genome quality while losing the epigenome. The *in silico* method, Modpolish, removed these modification-mediated errors without prior knowledge of modifications and uncovered the modified motifs while retaining the epigenome. When unknown modifications extensively shaped the genome, ONT with WGA or Modpolish produced nearly identical cgMLST profiles as hybrid ONT/Illumina did. On the other hand, the phylogeny of ONT-only genomes was disturbed by modification-mediated errors. Therefore, ONT with WGA or Modpolish is robust to modification-mediated errors without the need for additional Illumina sequencing.

### Quality reduction of ONT on novel modifications

Existing ONT basecalling algorithms only capture a few methylations (e.g., 5mC, 5hmc, 6mA) and ignore the vast amount of other modifications. Theoretically, species-specific modifications can be distinguished by training bespoke models for one organism (e.g., Taiyaki). But practically, it is infeasible to train models for hundreds of modifications in the biosphere. Especially in metagenomic sequencing, the usage of any particular model is biased against other modifications. For instance, a meta-epigenomic sequencing uncovered 22 methylation systems in a single microbial community [10]. Hence, if WGA is not an option, modification-mediated errors are better removed at the postassembly stage as each assembled contig can be polished independently.

### Limitations of ONT with WGA

The cost of WGA ONT is higher than ordinary sequencing due to several side effects of the amplification protocol. First, the amplified DNA may still contain a hyperbranched structure after Flap endonuclease (e.g., T7) cleavage. The hyperbranched DNA may block the pores during ONT sequencing and reduce the available pores and yields. In addition, the usage of endonuclease cleavage also decreased the read lengths. In conjunction with the uneven amplification, WGA requires higher coverage (∼100x) for reconstructing a complete genome than ordinary ONT sequencing (∼30x). Notably, the usage of WGA discards the entire methylome. The loss of modifications would prohibit any epigenetic study using ONT.

### Limitations of ONT with Modpolish

While Modpolish eliminated most modification-mediated errors, the correction power was lower in the ST1081 isolate. The lack of ST1081 genomes in NCBI RefSeq decreased the sensitivity of Modpolish. As the algorithm only corrects the loci of high evolutionary conservation, a sufficient number of closely-related genomes is necessary. Therefore, Modpolish is more suitable for common instead of rare lineages.

Nevertheless, Modpolish retains all modifications after ONT sequencing while WGA loses the epigenome. Epigenetic methylation has been thought to contribute to the rapid adaptation of resistance [11]. For instance, phase-variable adenine DNA methyltransferases (e.g., ModA11 and ModA12) increase susceptibility to cloxacillin and ciprofloxacin in *Neisseria meningitidis* [12]. The resistance due to overexpression of efflux pumps (e.g., sugE) has been linked to the lack of the Dcm-mediated 5mC silencing [13]. Therefore, Modpolish should be used when the epigenome is the focus of the study.

### Functional implications of the two modification systems

We discovered two pentanucleotide motifs, GCTGG (CCAGC) and GCAGC (GCTGC), specific to each of the two lineages (ST1081 and ST87). In ST1081, the GCTGG (CCAGC) motif is part of *chi* sites, hotspots of homologous recombination mediated by the RecBC enzyme [14, 15]. As phages cut by restriction enzymes are further degraded by RecBC [16], modifications on the GCTGG motif may be part of the defending system of ST1081, which protect itself against the RecBC cleavage.

In ST87 strains, the GCAGC/GCTGC (i.e., GCWGC) motif was the known target of the orphan methyltransferase M.BatI [17]. M.BatI produced fully-methylation on 5’
s-GCWGC-3′ and hemimethylation on 5′-GCSGC-3′. Reinvestigation of the modified sites in ST87 showed the existence of both GCWGC and GCSGC (Supplementary Figure S4). Interestingly, M.BatI increased toxicity when expressed in *E coli* in their study, which was concordant with the elevated virulence of ST87 strains.

Hence, the two lineages possessed two distinct modification systems for defensive purposes and increasing virulence. Although further investigations are required to assess their biological function, modifications that have acquired regulatory effects in bacteria are usually conservative within a clade [18]. Consequently, our *in silico* algorithm successfully utilize the conservation for correcting modification errors.

## Conclusion

This paper reported a set of unexpectedly low-quality genomes due to novel modifications untrained in the ONT basecalling model. The increasing number of new modifications found by single-molecular sequencing or high-resolution mass spectrometry will unavoidably reduce the ONT accuracy. New ONT flowcells, sequencing kits, and basecalling algorithms aim to resolve the homopolymer issue but not modification-mediated errors. Our study showed that these modification-mediated errors can be effectively corrected by preassembly amplification or postassembly polishing without additional short-read sequencing, producing high-quality genomes reliable for downstream analysis.

## Materials and Methods

### Bacterial isolates

Twelve *Listeria monocytogenes* isolates used in this study were obtained from hospitals recovered from listeriosis patients in Taiwan between 2019 and 2020. The isolates were submitted to the Taiwan Centers for Disease Control for further identification and genotyping. The isolates belonged to serogroups IIa (5 isolates), IIb (6 isolates), and IVb (1 isolate) and sequence type (ST) 1, ST5 (2 isolates), ST87 (4 isolates), ST101, ST155, ST378, ST1081, and ST1532.

### Whole genome sequencing

WGS of bacterial isolates was conducted in the Central Region laboratory of Taiwan CDC using the Illumina MiSeq sequencing platform (Illumina Co., USA) and the Nanopore sequencing platform (Oxford Nanopore Technologies, Inc., UK). DNA of bacterial isolates was extracted using the Qiagen DNeasy blood and tissue kit (Qiagen Co., Germany). Illumina DNA library construction was performed using the Illumina DNA Prep, (M) Tagmentation system (Illumina Co.), and sequencing was run with the MiSeq reagent kit version 3 (2X 300 bp), manipulated according to the manufacturer’s instructions. Nanopore DNA library construction was performed using the Rapid Barcoding Kit and sequencing was run using the MinION device and R9.4 chemistry.

### Removal of modifications of nucleotides using whole-genome amplification

DNA Bacterial Genomic DNA was amplified using the REPLI-g Advanced DNA Single Cell Kit (Qiagen, Hilden, Germany), manipulated according to the manufacturer’s instructions. The amplified DNA was purified using the KAPA HyperPure Beads (Roche, Basel, Switzerland) before subjecting to Nanopore sequencing.

### Assembly of sequence reads

Illumina sequence reads for each isolate were assembled using the SPAdes assembler version 3.12.0 (http://cab.spbu.ru/software/spades/) [19]; both Illumina sequence reads and Nanopore sequence reads for each isolates together were assembled to complete the full genomic sequences using the Unicycler Assembler [20]. The Nanopore reads for each isolate (in FAST5 file) were subjected to basecalling using the Guppy basecaller (https://nanoporetech.com/). In the ONT-only assembly, the sequences (in FASTQ file) were assembled using Flye (https://github.com/fenderglass/Flye)[21], then polished using the Racon (https://github.com/lbcb-sci/racon) [4], the Medaka (https://github.com/nanoporetech/medaka), and the Homopolish (https://github.com/ythuang0522/homopolish) [5]. Methylations (i.e., 5mC, 6mA) in the ONT-only genomes were called by Megalodon (https://github.com/nanoporetech/megalodon). The Integrative Genome Viewer (IGV) was used for visualizing the ONT modification errors [22]. The genome quality was assessed by fastmer (https://github.com/jts/assembly_accuracy).

### cgMLST analysis

Assembled Illumina contigs, assembled and polished Nanopore contigs, and assembled complete genomic sequence (obtained from assembling Illumina sequences and Nanopore sequences) for each isolate were used to generate core-gene multilocus sequence typing (cgMLST) profiles (based on 2,172 core genes) using an in-house-developed cgMLST profiling tool available on the cgMLST@Taiwan website (http://rdvd.cdc.gov.tw/cgMLST). Phylogenetic trees were constructed with cgMLST profiles using the minimum spanning tree algorithm and the tool provided on the cgMLST@Taiwan website.

### Overview of Modpolish

The proposed computational method, Modpolish, aims to remove modification-mediated errors by investigating the inconsistency of basecalled nucleotides, qualities of basecalled alleles, and evolutionary conservation at the modified loci. Modpolish is an extension from Homopolish, a polishing algorithm designed for correcting ONT homopolymer errors [5]. Figure 5 depicts the workflow of Modpolish. The closely-related genomes are first identified by screening against a compressed representation of microbial genomes. The genome sequences are then retrieved on the fly and compared with the draft genome. We only retain closely-related genomes of high nucleotide and structural similarity. Given the alignment matrix of reads, qualities, and homologs, Modpolish identifies potential-modified loci of inconsistent basecalling and low quality and only corrects the mismatch errors highly conserved in homologs. The details are described in the following sections.

**Figure 5.**
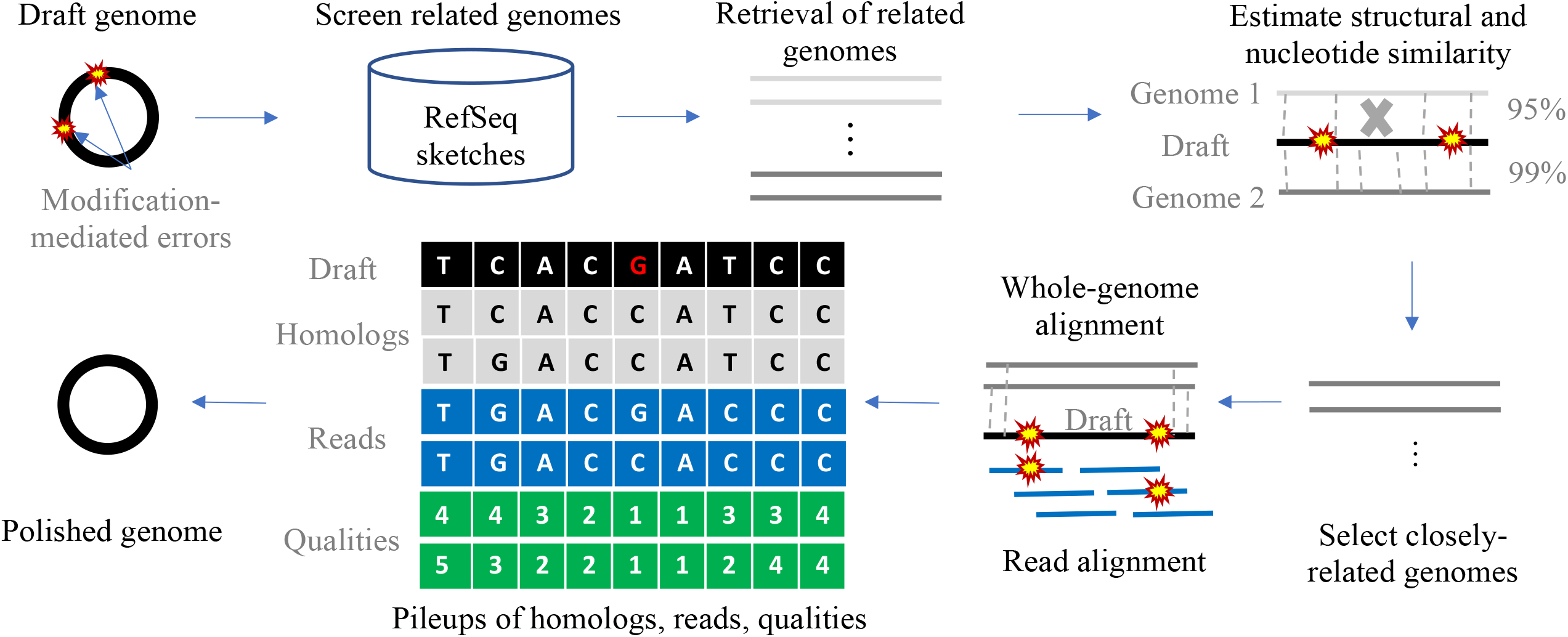
Illustration of Modpolish workflow. A set of closely-related genomes are first retrieved by screening the compressed sketches of RefSeq genomes. We retain the genomes with sufficient nucleotide and structural similarity. The selected genomes and ONT reads are aligned onto the draft genome, generating a pileup matrix of homologs, reads, and qualites. Modpolish only corrects modification-mediated errors with inconsistent read alleles, low quality, and high conservation in homologs.

### Collection of homologs by nucleotide and structural similarity

The draft genome (to be polished) is scanned against the virus, bacteria, or fungus genomes compressed by Mash as (MinHash) sketches, which is a reduced representation of all microbial genomes in NCBI RefSeq [23]. Subsequently, top *t* (default 20) closely related genomes will be retrieved on the fly. Mash estimated the Jaccard similarity between the draft and related genomes over a subset of *k*-mers. Though very fast, this method has low resolution at distinguishing closely-related genomes because the small subset of *k*-mers may not capture the few strain variations. Consequently, the genome similarity has to be re-estimated using more sensitive approaches.

Subsequently, each downloaded genome is compared against the draft genome using FastANI for computing the average nucleotide identity (ANI) at a higher resolution than Mash [24]. FastANI chops the two genomes into pieces and aligns them against each other for speedup. However, it only considers the aligned segments for ANI estimation and ignores the unaligned portions (Supplementary Figure S8(a)). The unaligned segments imply these two genomes differ by structural variations (i.e., vertically-/horizontally-transferred genes). As small and large variants are both genetic footprints of strain variations during evolution, Modpolish also computes the structural similarity (average-structural identity, ASI), defined as the percentage of aligned segments. We only retain the related genomes with sufficient ANI (>99%) and ASI (>90%) for subsequent error correction. These emprical cutoffs were determined by investigating the distributions of ANI and ASI in real microbial genomes.

### Correction of modification-mediated errors by reads and homologs

These closely-related genomes with sufficient ANI and ASI are aligned against the draft genome via minimap2 (with asm5 option) [25]. The raw ONT reads are also mapped against the draft genome by minimap2 (with map-ont option). We extract the basecalled nucleotides, basecalling qualities, and homologous alleles from the alignments. The aligned homologs, reads, and qualities are converted into a table of several summary statistics (Supplementary Figure S8(b)).

The summary statistics include the allele counts of A, T, C, and G separately for homologs and ONT reads, ignoring the insertion and deletion gaps. We identify the potentially modified sites according to the allele discordancy and average quality (see also Supplementary Figure S8(b)). The allele discordancy is the frequency of alternative alleles (i.e., non-major ones) at one locus. The average quality was computed by averaging the qscores from all read bases at the same locus. A potentially-modified locus is defined as the allele discordancy greater than 5% and the average quality score below 15, which were empirically observed from the modification-mediated errors.

For each potentially-modified locus, if all the homologous alleles are 100% conserved, we will correct the erroneous nucleotide into the alternative allele concordant with the homologs. These stringent criteria aimed for specificity instead of sensitivity, ensuring little or no false corrections would be made. We also implemented a motif-aware mode when the modification system is known in advance. If the user specifies a known modification motif (e.g., CCGAC), the program will additionally correct loci according to the provided pattern by lowering the homologous conservation ratio from 100% to 80%.

## Data and software availability

The genomes sequenced and assembled by Illumina, ONT, and WGA ONT are deposited in the NCBI with BioProject (xxxxxx). Modpolish was implemented as a subcommand in the Homopolish package, which is freely available at (https://github.com/ythuang0522/homopolish/).

## Supplementary Information

**Additional file 1**

Additional file 1 includes Supplementary Figures S1-8.

**Additional file 2**

Additional file 2 includes Supplementary Tables S1-10.

