## Supplementary Figure for "Correcting Modification-Mediated Errors in Nanopore Sequencing by Nucleotide Demodification and in silico Correction"

### Supplementary Figures

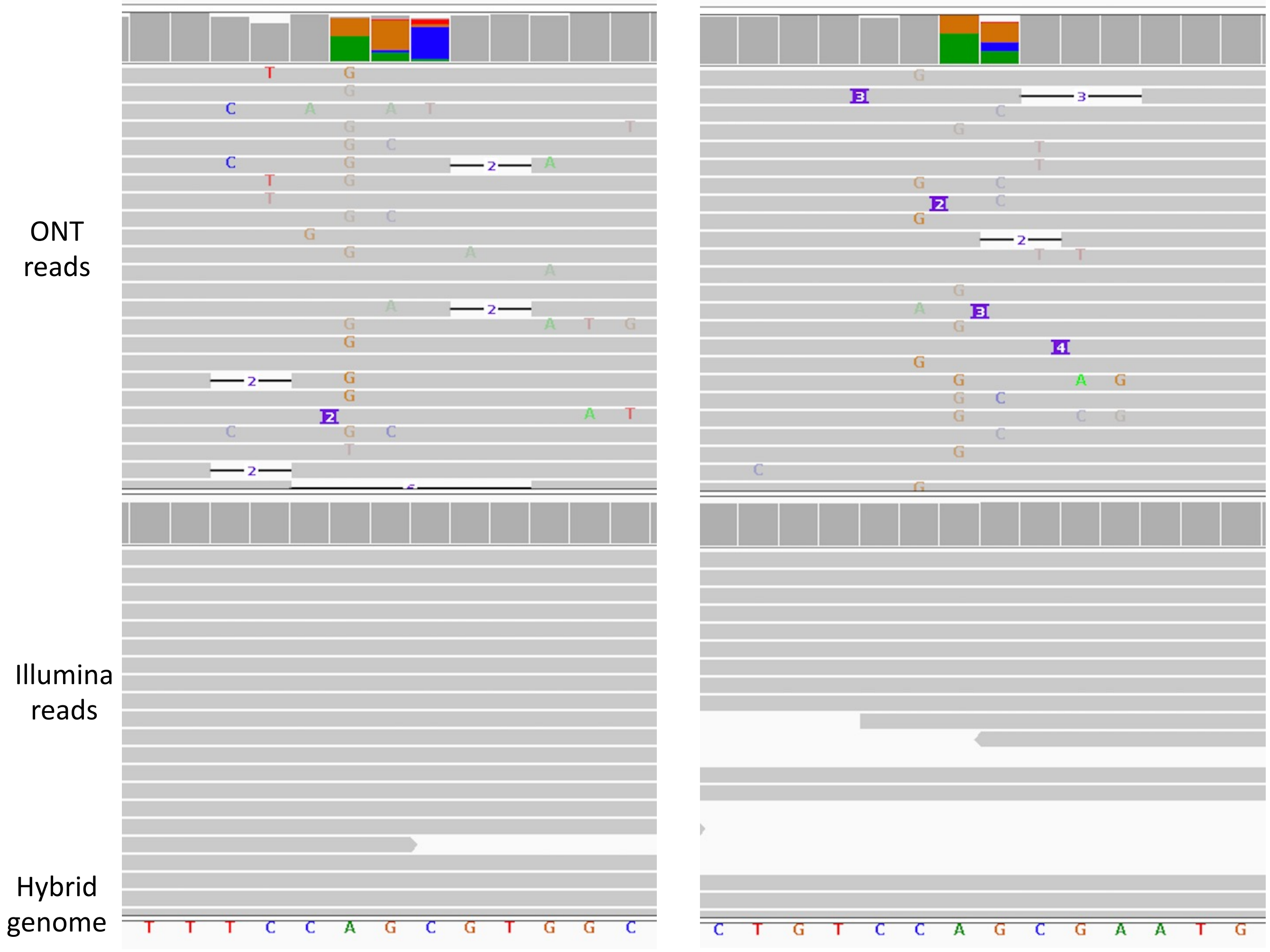

Supplementary Figure S1. Examples of ONT modification-mediated errors revealed by IGV. Top track: ONT reads; Middle track: Illumina reads; Bottom track: Hybrid assembled genome.

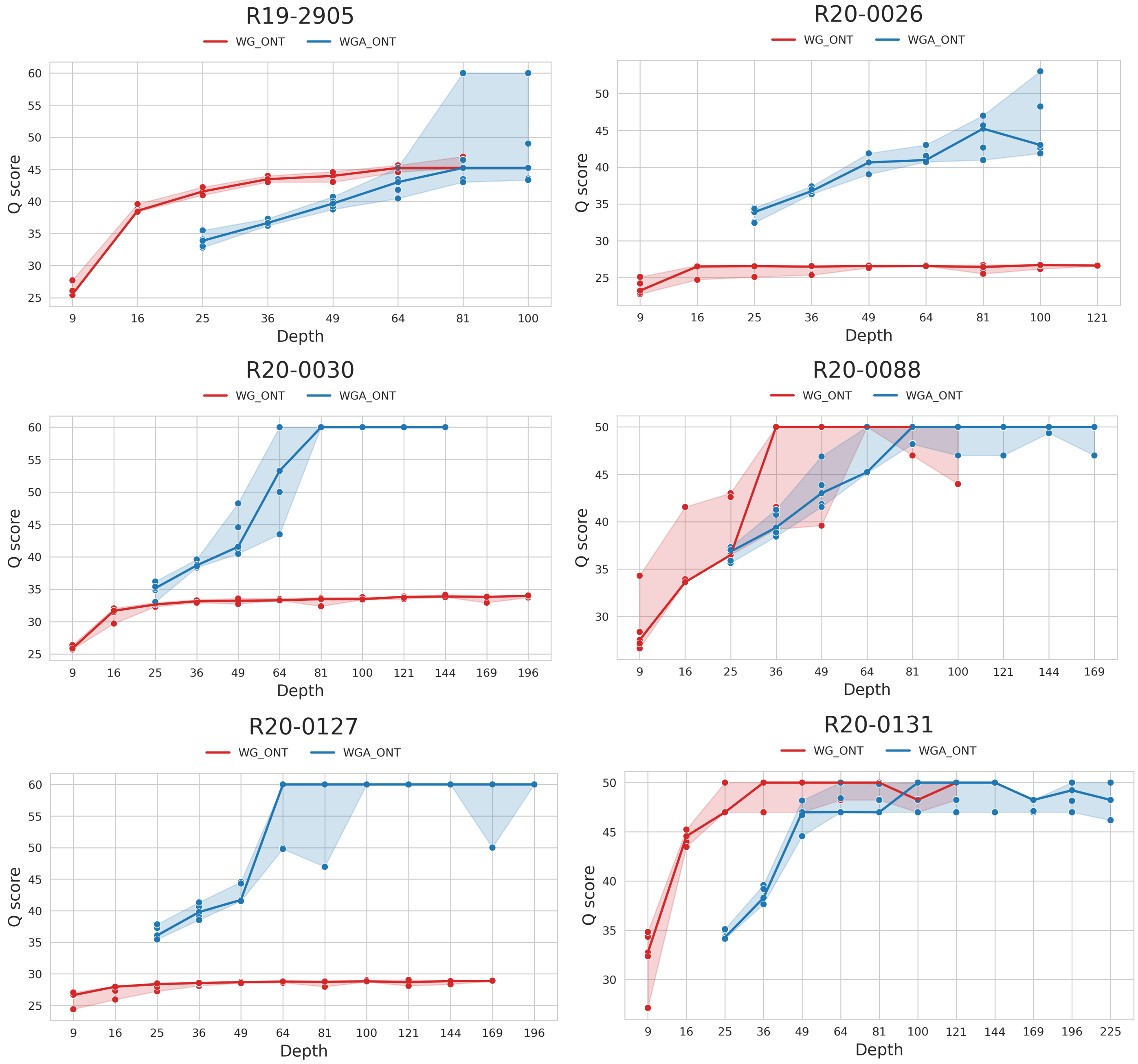

Supplementary Figure S2. Comparison of ONT-only and WGA-demodified ONT genome quality at different sequencing depth: R19-2904, R20-0026, R20-0030, R20-0088, R20-0127, and R20-0131.

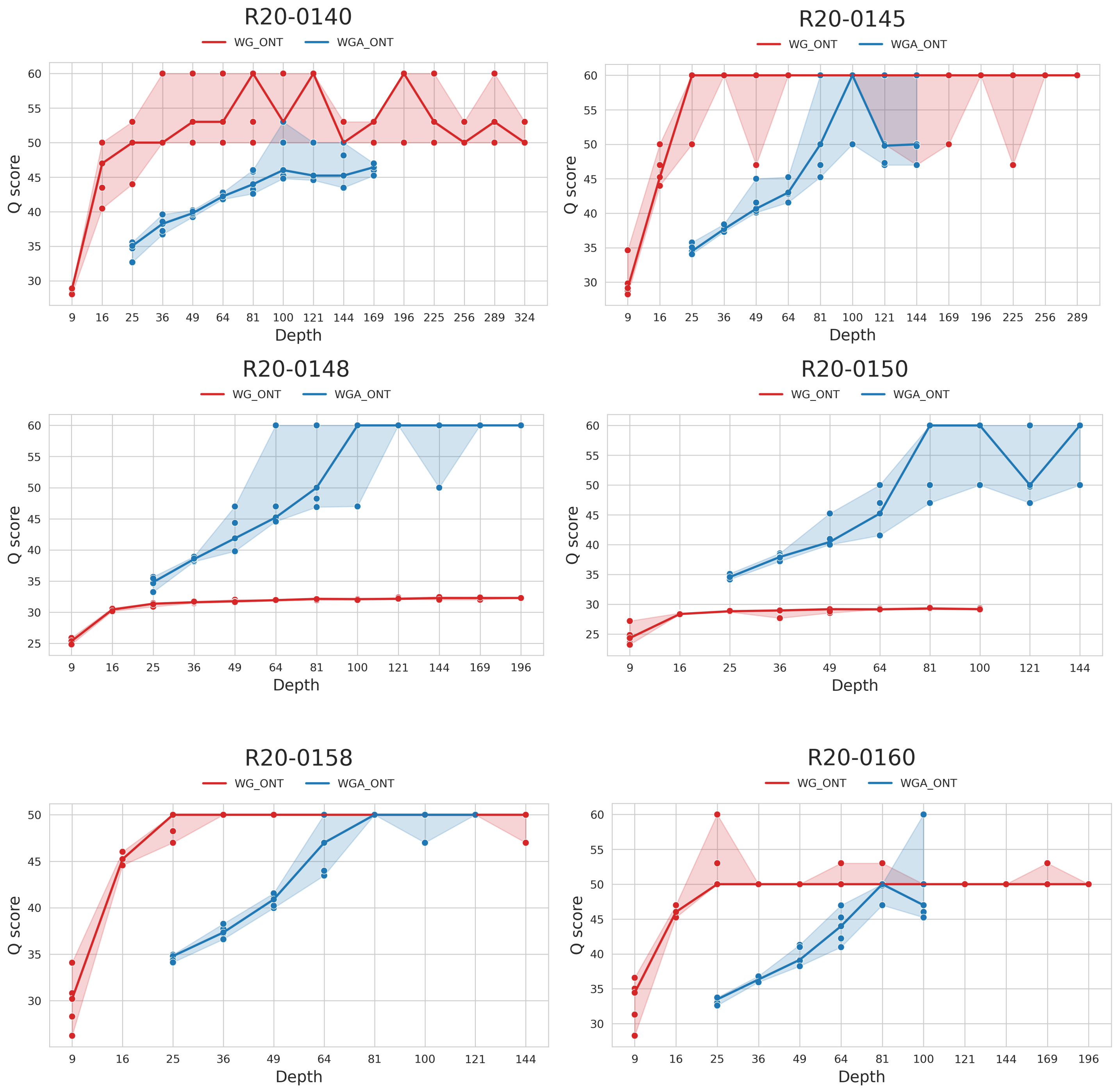

Supplementary Figure S3. Comparison of ONT-only and WGA-demodified ONT genome quality at different sequencing depth: R20-0140, R20-0145, R20-0148, R20-0150, R20-0158, and R20-0160.

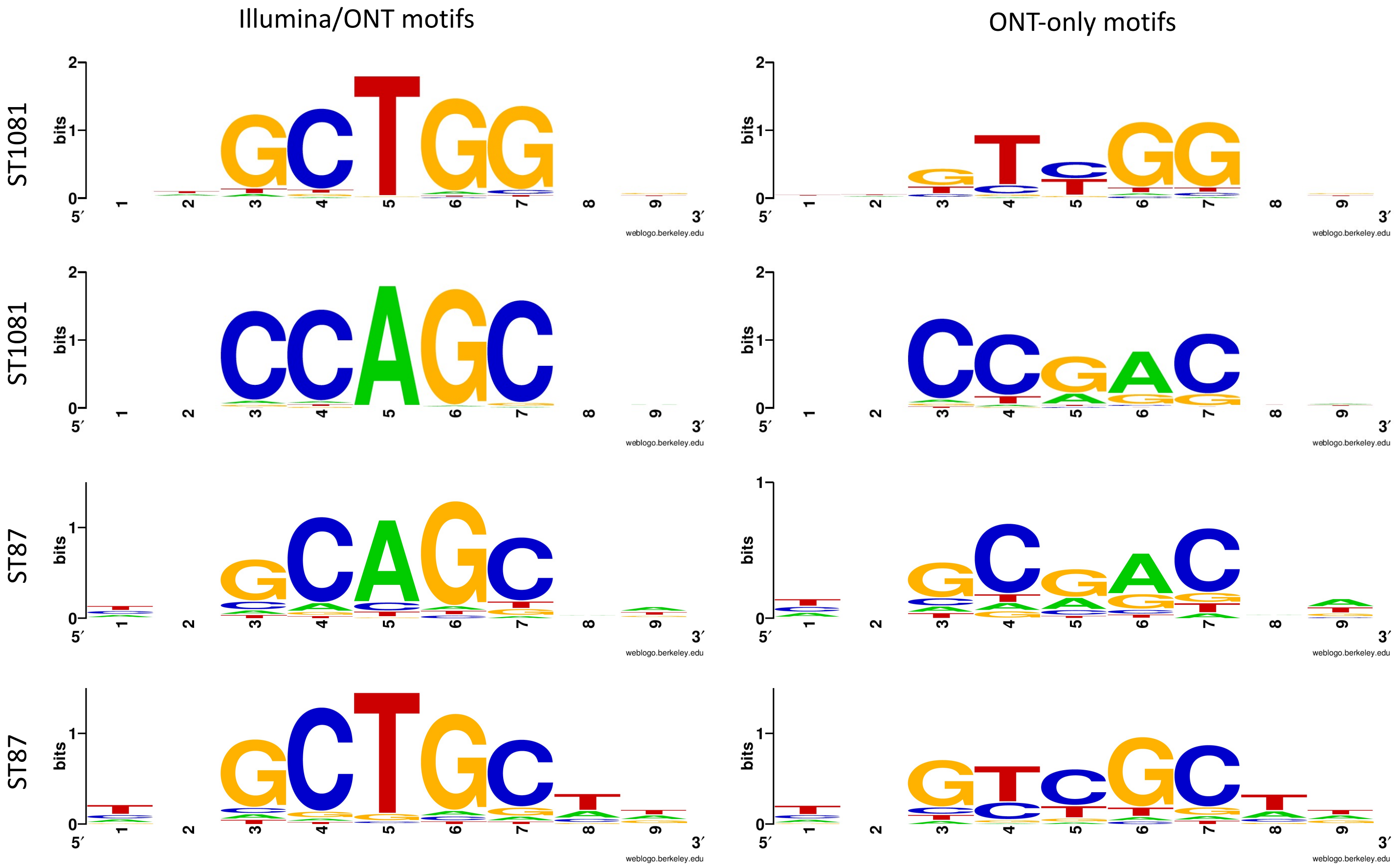

Supplementary Figure S4. Comparison of the modification motifs for the ST1081 and ST81 strains separately on the hybrid ONT/Illumina and ONT-only genomes.

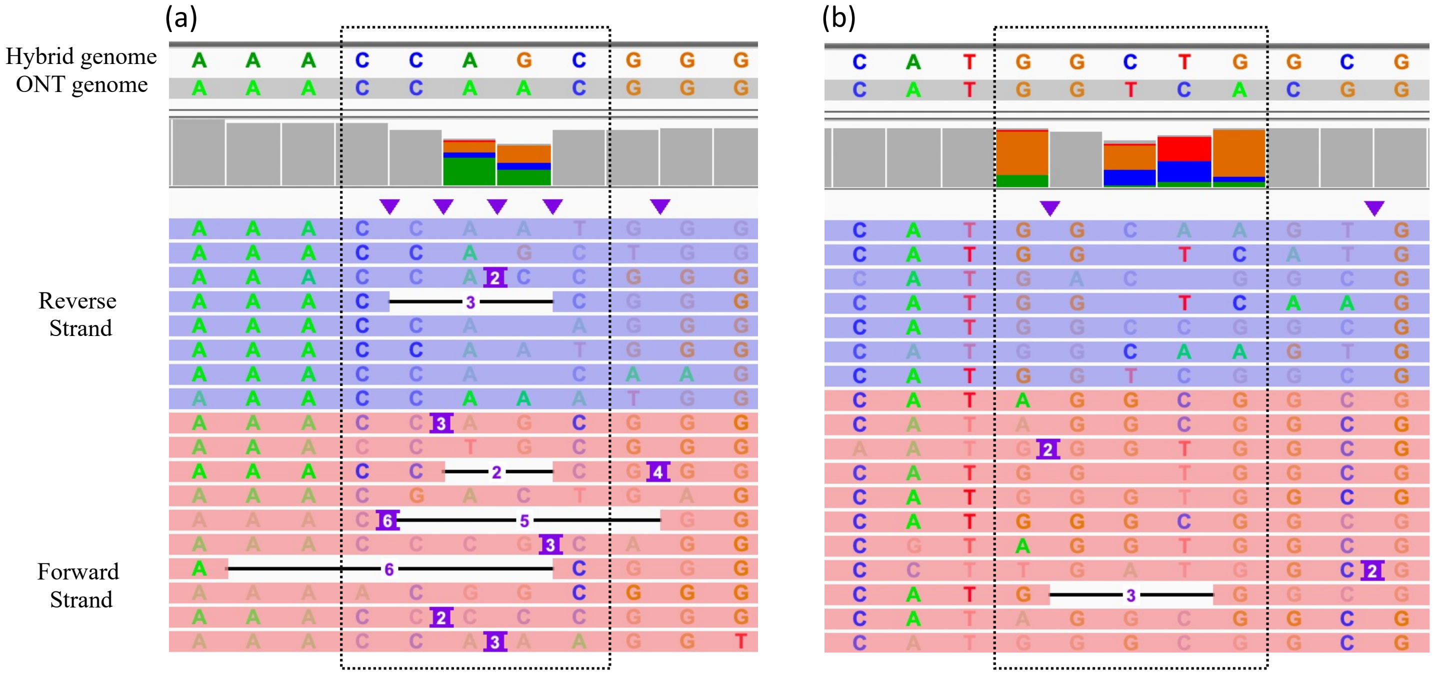

Supplementary Figure S5. Illustration of double-stranded modification by IGV. Basecalling errors were found on both the forward and reverse strands. (a) Sequencing errors for CCAGC motif; (b) sequencing errors for GGCTG motif.

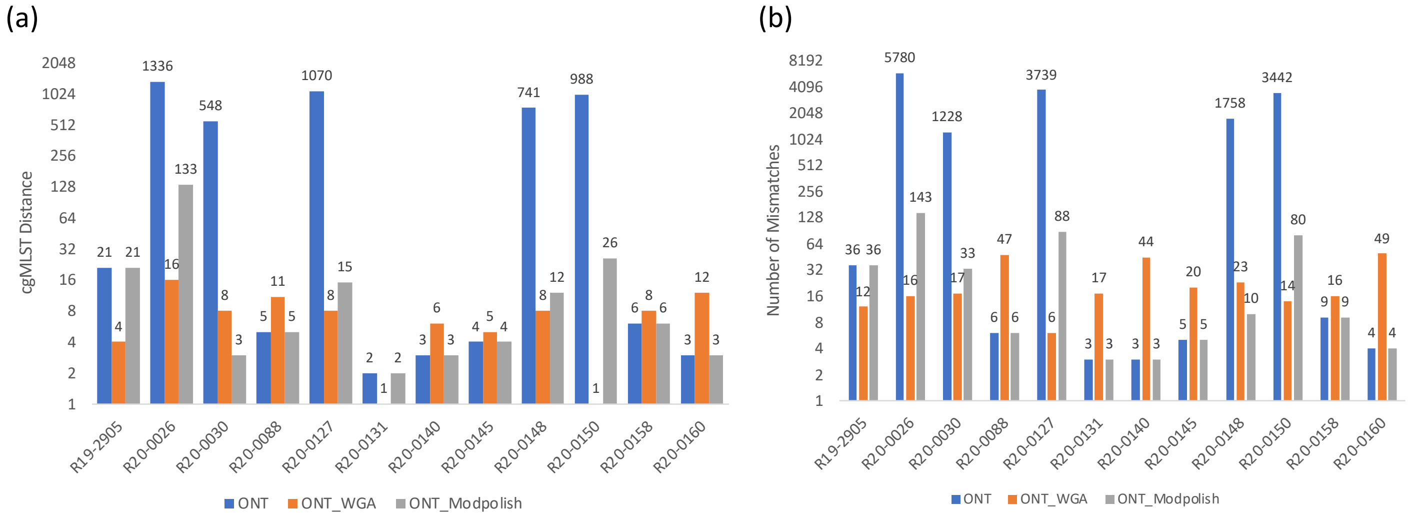

Supplementary Figure S6. Comparison of cgMLST distances and mismatches of ONT, WGA, and Modpolish with hybrid assemblies. (a) cgMLST distances; (b) numbers of mismatches.

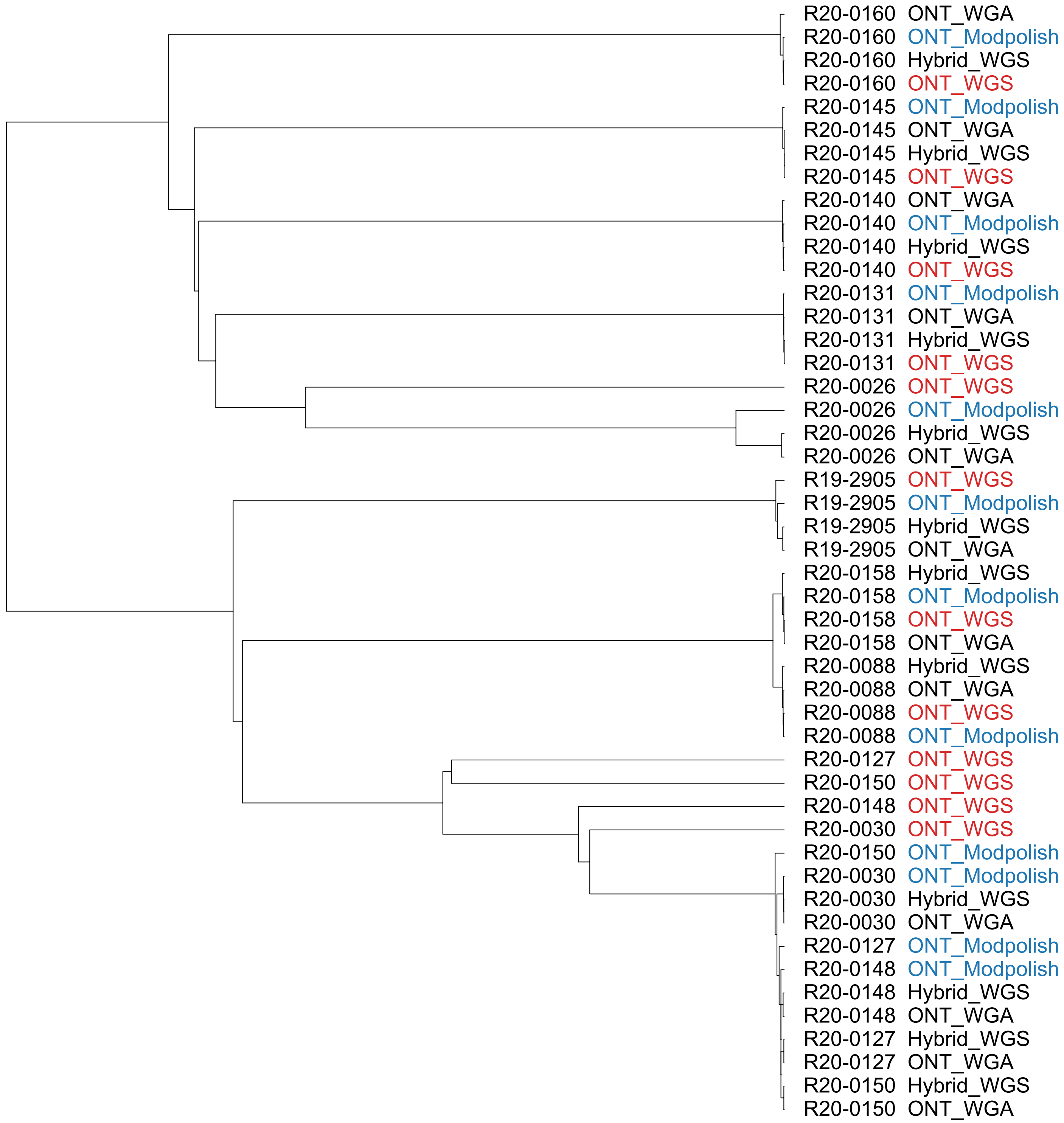

Supplementary Figure S7. The cgMLST phylogeny of the 12 *L monocytogene* strains. Each strain was ssembled by solely ONT (ONT_WGS), WGA-demodified ONT (ONT_WGA), ONT with Modpolish (ONT_Modpolish), and hybrid ONT/Illumina genomes (Hybrid_WGS).

(a)

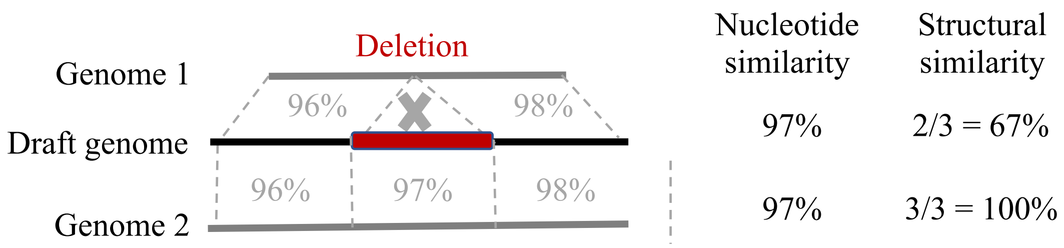

(b)

| | | | | | | | | | | | |
|---|---|---|---|---|---|---|---|---|---|---|---|
| True genome | | A | G | C | T | G | **C** | **G** | T | G | G |
| Draft genome | | A | G | C | T | G | **T** | **T** | T | G | G |
| Homologous alleles | A | 20 | | | 10 | | | | | | 1 |
| | C | | | 20 | | 10 | 20 | | | | |
| | G | | 20 | | | 10 | | 20 | | 20 | 19 |
| | T | | | | 10 | | | | 20 | | |
| Read alleles | A | 25 | | | 2 | 1 | 2 | 1 | 1 | | |
| | C | | | 23 | | 1 | 1 | 0 | | | |
| | G | | 25 | | | 1 | 0 | 2 | | 20 | 20 |
| | T | | | | 18 | 17 | 17 | 17 | 19 | | |
| Average quality | | 20 | 20 | 20 | 16 | 12 | 8 | 12 | 16 | 20 | 20 |
| Allele discordance | | 0 | 0 | 0 | 10 | 15 | **15** | **15** | 5 | 0 | 0 |
| Homolog conservation | | 100 | 100 | 100 | 50 | 50 | **100** | **100** | 100 | 100 | 95 |

Supplementary Figure S8. Similarity computation and allele count statistics. (a) Comparision of structural and nucleotide similarities; (b) Illustration of homologous and read alleles pileup.
