## Supplementary Table for "Correcting Modification-Mediated Errors in Nanopore Sequencing by Nucleotide Demodification and in silico Correction"

### Supplementary Tables

#### Table of Contents

Supplementary Table S1. Illumina sequencing statistics of the 12 samples. 2
Supplementary Table S2. ONT sequencing statistics of the 12 samples. 2
Supplementary Table S3. Assembly statistics of ONT-only sequencing 2
Supplementary Table S4. Assembly statistics of hybrid ONT/Illumina sequencing 3
Supplementary Table S5. Quality assessment of ONT-only genomes polished by Medaka and Homopolish. 3
Supplementary Table S6. Numbers of 5mC and 6mA modifications identified by Megalodon. 3
Supplementary Table S7. ONT sequencing statistics of the 12 WGA-demodified samples. 4
Supplementary Table S8. Assembly statistics of WGA-demodified ONT sequencing 4
Supplementary Table S9. Quality assessment of WGA-demodified ONT genomes 4
Supplementary Table S10. Quality assessment of ONT genomes corrected by Modpolish. 5

Supplementary Table S1. Illumina sequencing statistics of the 12 samples.

| | Read Number | Max Length | N50 Length | Total Bases |
|---|---|---|---|---|
| **R19-2905** | 935808 | 301 | 301 | 251516220 |
| **R20-0026** | 681360 | 301 | 301 | 199375540 |
| **R20-0030** | 765356 | 301 | 301 | 223843041 |
| **R20-0088** | 792890 | 300 | 300 | 211480191 |
| **R20-0127** | 1143744 | 301 | 301 | 310918904 |
| **R20-0131** | 808834 | 300 | 300 | 217882330 |
| **R20-0140** | 1084336 | 301 | 301 | 296601904 |
| **R20-0145** | 1046572 | 301 | 301 | 283615978 |
| **R20-0148** | 1340102 | 300 | 300 | 359675819 |
| **R20-0150** | 1153082 | 301 | 301 | 310089709 |
| **R20-0158** | 1254996 | 301 | 301 | 343552792 |
| **R20-0160** | 1364554 | 301 | 301 | 370035262 |

Supplementary Table S2. ONT sequencing statistics of the 12 samples.

| | Read Number | Max Length | N50 Length | Total Bases |
|---|---|---|---|---|
| **R19-2905** | 103489 | 113264 | 6988 | 311272973 |
| **R20-0026** | 109179 | 133103 | 8776 | 409731944 |
| **R20-0030** | 318211 | 109539 | 4237 | 658332239 |
| **R20-0088** | 95611 | 110293 | 9450 | 345134126 |
| **R20-0127** | 136705 | 113937 | 8906 | 552032094 |
| **R20-0131** | 100549 | 108202 | 8433 | 381762248 |
| **R20-0140** | 377099 | 74028 | 5124 | 990811868 |
| **R20-0145** | 291672 | 180955 | 5879 | 877310303 |
| **R20-0148** | 265358 | 80144 | 4282 | 589230943 |
| **R20-0150** | 90990 | 107273 | 9529 | 364792267 |
| **R20-0158** | 111136 | 127984 | 9827 | 438704763 |
| **R20-0160** | 124026 | 106338 | 10960 | 593804240 |

Supplementary Table S3. Assembly statistics of ONT-only sequencing

| | Contig Number | Max Length | N50 Length | Total Bases |
|---|---|---|---|---|
| **R19-2905** | 5 | 2408977 | 2408977 | 3139752 |
| **R20-0026** | 2 | 2941441 | 2941441 | 2947228 |
| **R20-0030** | 2 | 2953610 | 2953610 | 2958812 |
| **R20-0088** | 3 | 2949122 | 2949122 | 3074902 |
| **R20-0127** | 2 | 2992895 | 2992895 | 2998908 |
| **R20-0131** | 2 | 2952984 | 2952984 | 2958875 |
| **R20-0140** | 4 | 1802146 | 1802146 | 2932180 |
| **R20-0145** | 2 | 2895673 | 2895673 | 2901698 |
| **R20-0148** | 2 | 2912509 | 2912509 | 2917601 |
| **R20-0150** | 4 | 3015129 | 3015129 | 3070241 |
| **R20-0158** | 2 | 2945198 | 2945198 | 2951211 |
| **R20-0160** | 2 | 2986867 | 2986867 | 2992875 |

Supplementary Table S4. Assembly statistics of hybrid ONT/Illumina sequencing

| | Contig Number | Max Length | N50 Length | Total Bases |
|---|---|---|---|---|
| **R19-2905** | 1 | 3160104 | 3160104 | 3160104 |
| **R20-0026** | 1 | 2941412 | 2941412 | 2941412 |
| **R20-0030** | 1 | 2953630 | 2953630 | 2953630 |
| **R20-0088** | 2 | 2949137 | 2949137 | 3009037 |
| **R20-0127** | 1 | 2992871 | 2992871 | 2992871 |
| **R20-0131** | 1 | 2953003 | 2953003 | 2953003 |
| **R20-0140** | 1 | 2953885 | 2953885 | 2953885 |
| **R20-0145** | 1 | 2895698 | 2895698 | 2895698 |
| **R20-0148** | 1 | 2912653 | 2912653 | 2912653 |
| **R20-0150** | 1 | 3015149 | 3015149 | 3015149 |
| **R20-0158** | 1 | 2945208 | 2945208 | 2945208 |
| **R20-0160** | 1 | 2986889 | 2986889 | 2986889 |

Supplementary Table S5. Quality assessment of ONT-only genomes polished by Medaka and Homopolish.

| | **Mismatch** | **insertion** | **deletion** | **Q score** |
|---|---|---|---|---|
| **R19-2905** | 36 | 20 | 23 | 47 |
| **R20-0026** | 5780 | 304 | 166 | 27 |
| **R20-0030** | 1228 | 10 | 17 | 34 |
| **R20-0088** | 6 | 15 | 16 | 50 |
| **R20-0127** | 3739 | 64 | 20 | 29 |
| **R20-0131** | 3 | 12 | 24 | 50 |
| **R20-0140** | 3 | 14 | 16 | 60 |
| **R20-0145** | 5 | 7 | 18 | 60 |
| **R20-0148** | 1758 | 15 | 7 | 32 |
| **R20-0150** | 3442 | 62 | 62 | 29 |
| **R20-0158** | 9 | 11 | 28 | 50 |
| **R20-0160** | 4 | 7 | 10 | 60 |

Supplementary Table S6. Numbers of 5mC and 6mA modifications identified by Megalodon.

| | **5mC** | **6mA** |
|---|---|---|
| **R19-2905** | 343754 | 189433 |
| **R20-0026** | 295057 | 142989 |
| **R20-0030** | 339836 | 218745 |
| **R20-0088** | 218745 | 155818 |
| **R20-0127** | 268977 | 114577 |
| **R20-0131** | 263553 | 141580 |
| **R20-0140** | 324727 | 223724 |
| **R20-0145** | 336799 | 196019 |
| **R20-0148** | 245296 | 98068 |
| **R20-0150** | 304172 | 147307 |
| **R20-0158** | 262102 | 139438 |
| **R20-0160** | 260236 | 126020 |

Supplementary Table S7. ONT sequencing statistics of the 12 WGA-demodified samples.

| | Read Number | Max Length | N50 Length | Total Bases |
|---|---|---|---|---|
| **R19-2905** | 188066 | 68252 | 4008 | 381295211 |
| **R20-0026** | 161346 | 74725 | 3998 | 323072380 |
| **R20-0030** | 225282 | 80954 | 3778 | 429041543 |
| **R20-0088** | 295820 | 98758 | 3760 | 551208279 |
| **R20-0127** | 351965 | 105728 | 3754 | 660699365 |
| **R20-0131** | 361034 | 94318 | 4085 | 744673304 |
| **R20-0140** | 320358 | 90561 | 3452 | 564265875 |
| **R20-0145** | 227362 | 64678 | 3783 | 431307951 |
| **R20-0148** | 342611 | 75934 | 3653 | 633592810 |
| **R20-0150** | 224242 | 57070 | 3896 | 443942733 |
| **R20-0158** | 218499 | 73549 | 3798 | 398753574 |
| **R20-0160** | 183849 | 90537 | 3810 | 338761482 |

Supplementary Table S8. Assembly statistics of WGA-demodified ONT sequencing

| | Contig Number | Max Length | N50 Length | Total Bases |
|---|---|---|---|---|
| **R19-2905** | 6 | 2408926 | 2408926 | 3163109 |
| **R20-0026** | 3 | 1498066 | 1498066 | 2947083 |
| **R20-0030** | 6 | 1375174 | 1368605 | 2970480 |
| **R20-0088** | 3 | 2460960 | 2460960 | 2966134 |
| **R20-0127** | 4 | 2303825 | 2303825 | 3011536 |
| **R20-0131** | 2 | 1519287 | 1519287 | 2946950 |
| **R20-0140** | 8 | 1104249 | 942128 | 2968716 |
| **R20-0145** | 5 | 1214009 | 917883 | 2907726 |
| **R20-0148** | 5 | 1585007 | 1585007 | 2944803 |
| **R20-0150** | 3 | 1555559 | 1555559 | 2981832 |
| **R20-0158** | 1 | 2974086 | 2974086 | 2974086 |
| **R20-0160** | 7 | 1290018 | 1264214 | 3038421 |

Supplementary Table S9. Quality assessment of WGA-demodified ONT genomes

| | **Mismatch** | **insertion** | **deletion** | **Q score** |
|---|---|---|---|---|
| **R19-2905** | 12 | 281 | 23 | 90 |
| **R20-0026** | 16 | 6456 | 30 | 53 |
| **R20-0030** | 17 | 10 | 6033 | 90 |
| **R20-0088** | 47 | 35 | 2065 | 50 |
| **R20-0127** | 6 | 17 | 35 | 90 |
| **R20-0131** | 17 | 40 | 31 | 50 |
| **R20-0140** | 44 | 2630 | 27 | 53 |
| **R20-0145** | 20 | 10 | 17 | 90 |
| **R20-0148** | 23 | 7917 | 23 | 90 |
| **R20-0150** | 14 | 14 | 6023 | 90 |
| **R20-0158** | 16 | 19 | 16 | 50 |
| **R20-0160** | 49 | 30 | 42 | 90 |

Supplementary Table S10. Quality assessment of ONT genomes corrected by Modpolish.

| | Mismatch | insertion | deletion | Q score |
|---|---|---|---|---|
| **R19-2905** | 36 | 20 | 23 | 46 |
| **R20-0026** | 143 | 44 | 24 | 45 |
| **R20-0030** | 33 | 3 | 17 | 60 |
| **R20-0088** | 6 | 15 | 16 | 50 |
| **R20-0127** | 88 | 64 | 20 | 50 |
| **R20-0131** | 3 | 12 | 24 | 50 |
| **R20-0140** | 3 | 14 | 16 | 60 |
| **R20-0145** | 5 | 5 | 18 | 60 |
| **R20-0148** | 10 | 5 | 7 | 60 |
| **R20-0150** | 80 | 62 | 13 | 50 |
| **R20-0158** | 9 | 11 | 28 | 50 |
| **R20-0160** | 4 | 7 | 10 | 60 |
